# The Evolution of Fluoroquinolone-Resistance in *Mycobacterium tuberculosis* is Modulated by the Genetic Background

**DOI:** 10.1101/659045

**Authors:** Rhastin A. D. Castro, Amanda Ross, Lujeko Kamwela, Miriam Reinhard, Chloé Loiseau, Julia Feldmann, Sonia Borrell, Andrej Trauner, Sebastien Gagneux

## Abstract

Fluoroquinolones (FQ) form the backbone in experimental treatment regimens against drug-susceptible tuberculosis. However, little is known on whether the genetic variation present in natural populations of *Mycobacterium tuberculosis* (*Mtb*) affects the evolution of FQ-resistance (FQ-R). To investigate this question, we used a set of *Mtb* strains that included nine genetically distinct drug-susceptible clinical isolates, and measured their frequency of resistance to the FQ ofloxacin (OFX) *in vitro*. We found that the *Mtb* genetic background led to differences in the frequency of OFX-resistance (OFX-R) that spanned two orders of magnitude and substantially modulated the observed mutational profiles for OFX-R. Further *in vitro* assays showed that the genetic background also influenced the minimum inhibitory concentration and the fitness effect conferred by a given OFX-R mutation. To test the clinical relevance of our *in vitro* work, we surveyed the mutational profile for FQ-R in publicly available genomic sequences from clinical *Mtb* isolates, and found substantial *Mtb* lineage-dependent variability. Comparison of the clinical and the *in vitro* mutational profiles for FQ-R showed that 45% and 19% of the variability in the clinical frequency of FQ-R *gyrA* mutations in Lineage 2 and Lineage 4 strains, respectively, can be attributed to how *Mtb* evolves FQ-R *in vitro*. As the *Mtb* genetic background strongly influenced the evolution of FQ-R *in vitro*, we conclude that the genetic background of *Mtb* also impacts the evolution of FQ-R in the clinic.

**Significance:** Newer generations of fluoroquinolones form the backbone in many experimental treatment regimens against *M. tuberculosis* (*Mtb*). While the genetic variation in natural populations of *Mtb* can influence resistance evolution to multiple different antibiotics, it is unclear whether it modulates fluoroquinolone-resistance evolution as well. Using a combination of *in vitro* assays coupled with genomic analysis of clinical isolates, we provide the first evidence illustrating the *Mtb* genetic background’s substantial role in fluoroquinolone-resistance evolution, and highlight the importance of bacterial genetics when studying the prevalence of fluoroquinolone-resistance in *Mtb*. Our work may provide insights into how to maximize the timespan in which fluoroquinolones remain effective in clinical settings, whether as part of current standardized regimens, or in new regimens against *Mtb*.

## Introduction

Antimicrobial resistance (AMR) poses a major threat to our ability to treat infectious diseases (1, 2). The rise of AMR is a complex phenomenon with a broad range of contributing socioeconomic and behavioural factors (3–7). However, the emergence of AMR within any pathogen population is ultimately an evolutionary process (8, 9). This evolutionary process is influenced by multiple factors, including drug pressure and pathogen genetics. Firstly, the drug type and drug concentration can affect the type and relative frequencies of AMR mutations observed in a given pathogen population (also known as the mutational profile for AMR) (9–14). Secondly, pathogen populations comprise genetically distinct strains, and this genetic variation may also influence AMR evolution (15–17). Different pathogen genetic backgrounds can have different baseline susceptibilities to a given drug (18, 19), which consequently can affect patient treatment outcomes (20). The genetic background has also been shown to modulate the acquisition and prevalence of AMR (11, 15, 21, 22), the mutational profile for AMR (11, 15, 16, 23), and the phenotypic effects of AMR mutations (24–28). Studying the interplay between pathogen genetics and drug pressure is therefore important in understanding how to restrict the prevalence of AMR in pathogen populations.

AMR in *Mycobacterium tuberculosis* (*Mtb*), the aetiological agent of human tuberculosis (TB), is of particular importance. *Mtb* infections globally cause the highest rate of mortality due to a single infectious agent both in general, and due to AMR specifically (29). Although the genetic variation in *Mtb* is small compared to other bacterial pathogens (17, 30), several studies have shown that this limited genetic variation influences AMR phenotypes and prevalence (15, 17, 24, 28, 31). The global genetic diversity of *Mtb* comprises seven phylogenetic lineages (17, 30), and *Mtb* strains belonging to the Lineage 2 Beijing/W genetic background have repeatedly been associated with multidrug-resistant TB (MDR-TB; defined as an infection from an *Mtb* strain that is resistant to at least isoniazid and rifampicin) both *in vitro* and in clinical settings (4, 11, 15, 21, 22).

One strategy to reduce the emergence of AMR in *Mtb* is the development of new, shorter treatment regimens (32, 33). Many such experimental regimens use third- or fourth-generation fluoroquinolones (FQ) against drug-susceptible *Mtb* (32–36). However, FQs have long been integral to treating MDR-TB (37), and the previous use of FQ has led to the emergence of FQ-resistance (FQ-R) in clinical strains of *Mtb* (7, 38–40). FQ-R is one of the defining properties of extensively drug-resistant TB (XDR-TB), and XDR-TB accounts for 8.5% of MDR-TB cases (29). Understanding how FQ-R is acquired in natural populations of *Mtb* may allow for the development of tools or strategies to mitigate further increases in FQ-R prevalence.

In *Mtb*, the sole target of FQ is DNA gyrase (10, 38, 41–43). Consequently, clinically relevant FQ-R in *Mtb* is primarily due to a limited set of chromosomal mutations located within the “quinolone-resistance-determining region” (QRDR) of the *gyrA* and *gyrB* genes, which encode DNA gyrase (22, 38, 39). No horizontal gene-transfer (HGT) or plasmid-based resistance to FQ has been documented in *Mtb* (44, 45). Studying FQ-R evolution in *Mtb* populations thus provides a promising setting for elucidating how the genetic background may affect the emergence and maintenance of clinically relevant chromosomal AMR mutations in bacterial populations.

While a great deal of literature exists on the biochemical mechanisms leading to the FQ-R phenotype in *Mtb* (10, 41–43, 46, 47), little is known on the evolutionary dynamics of FQ-R in different populations of *Mtb*. Given that antimicrobial regimens against *Mtb* infections use standardized, empirical dosing strategies (29), it is unclear whether different *Mtb* genetic backgrounds would acquire FQ-R at the same frequency when exposed to the same antimicrobial concentration. Whether the *Mtb* genetic background would also modulate the mutational profile for FQ-R, or the phenotypic effects of FQ-R mutations, is unknown. Such knowledge may provide insights on how to maintain or prolong the efficiency of FQs against different genetic variants of *Mtb* in the clinic.

In this study, we tested whether the *Mtb* genetic background plays a role in the evolution of FQ-R. Specifically, we showed that the *Mtb* genetic background can lead to differences in the frequency of FQ-R emergence that span two-orders of magnitude, as well as substantially modulate the mutational profile for FQ-R. We further demonstrated that the phenotypic effects of clinically relevant FQ-R mutations differed depending on the *Mtb* genetic background they were present in. Analysis of publicly available genomic sequences from clinical *Mtb* isolates also revealed a positive association between the FQ-R mutational profiles observed *in vitro* and the mutational profiles observed in the clinic. Taken together, we showed that the *Mtb* genetic background had a considerable role in evolution of FQ-R in the clinic.

## Results

### Frequency of ofloxacin-resistance in *M. tuberculosis* is strain-dependent

We first tested for whether the *Mtb* genetic background led to differences in the frequency of FQ-R acquisition. To do so, we performed a Luria-Delbrück fluctuation analysis on nine drug-susceptible and genetically distinct *Mtb* clinical strains belonging to Lineage 1 (L1), Lineage 2 (L2) and Lineage 4 (L4) (See SI Appendix, Table S1) (17, 30, 48, 49). We measured their frequency of resistance *in vitro* to the FQ ofloxacin (OFX), as OFX was used extensively to treat MDR-TB patients in the past. Given that anti-TB treatment regimens use standardized drug concentrations (29), we also measured the frequency of resistance to the same concentration of OFX (4 µg/mL) for all nine strains. We observed significant strain-dependent variation in the frequency of OFX-resistance (OFX-R) at 4 µg/mL, with the difference spanning two orders of magnitude (Fig. 1A; *P* = 2.2 × 10^-16^, Kruskal-Wallis). Several of the nine drug-susceptible *Mtb* strains contained missense substitutions in DNA gyrase that are not associated with FQ-R (See SI Appendix, Table S2) (49). These mutations are phylogenetic markers that reflect the population structure of *Mtb* and cannot be avoided if strains from different *Mtb* lineages are used (17, 30). We found no evidence for any associations between the presence a given phylogenetic DNA gyrase missense mutation and the frequency of OFX-R acquired.

**Fig. 1.**
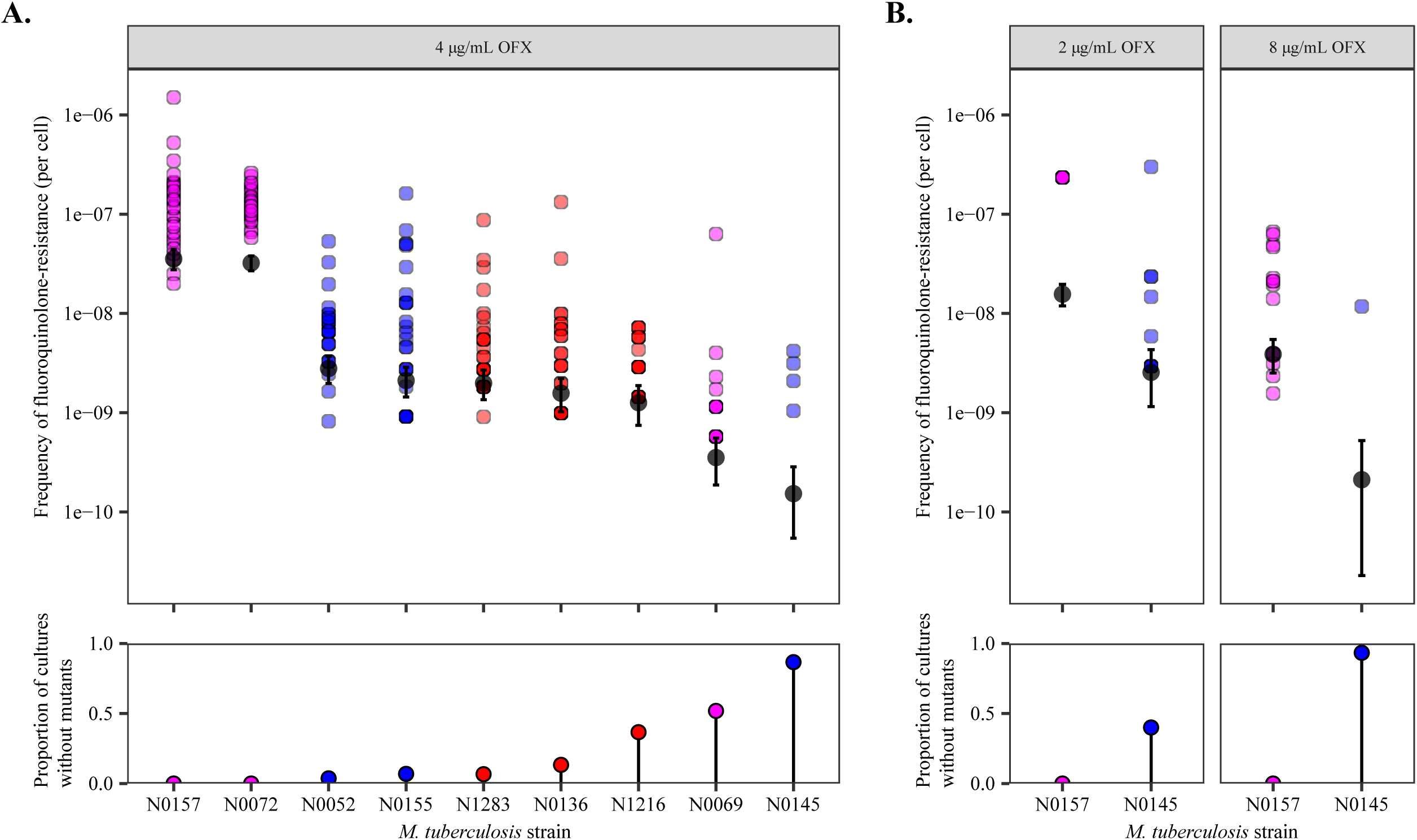
Variation in the frequency of ofloxacin-resistance between genetically distinct, wild-type *M. tuberculosis* strains. **A.** Frequency of ofloxacin-resistance at 4 µg/mL ofloxacin (OFX), as measured by fluctuation analysis. Top panel: Coloured points represent the frequency of resistant mutants per cell per parallel culture, with darker points representing multiple cultures with the same frequency. Colours denote the lineage that the *M. tuberculosis* strain belongs to (L1 = pink; L2 = blue; L4 = red). Grey points represent the estimated number of mutations per cell per strain as calculated by MSS-MLE, while black bars denote the respective 95% confidence intervals. Bottom panel: the percentage of parallel cultures lacking OFX-resistant mutants. **B**. Frequency of ofloxacin-resistance at 2 or 8 µg/mL OFX.

The concentration of the antimicrobial can affect the observed frequencies of AMR in *Mtb* (10, 11, 13). Therefore, we tested whether changing the selective concentration of OFX would affect the relative differences in strain-specific OFX-R frequencies. For the sake of simplicity, we tested only two strains, with each strain at the opposite extremes of the frequency of resistance to 4 µg/mL OFX, as shown in Fig. 1A: N0157 (high OFX-R frequency) and N0145 (low OFX-R frequency). We found that the frequency of OFX-R remained one to two-orders of magnitude higher in N0157 than in N0145 across all the concentrations we tested (Fig. 1B, *P* = 2.46 x 10^-5^ for 2 µg/mL OFX, and *P* = 4.03 x 10^-6^ for 8 µg/mL OFX, Wilcoxon rank-sum test). The N0157 strain had nearly confluent growth at 2 µg/mL OFX, which is the OFX concentration that has been shown to inhibit 95% of *Mtb* strains that have not been previously exposed to OFX, but does not inhibit *Mtb* strains that are considered resistant to OFX in the clinic (18, 19). This suggested that N0157 has low-level resistance to OFX, despite having no mutation in the QRDR. Meanwhile, at 8 µg/mL OFX, we observed only four resistant colonies for N0145 across all samples, with all colonies arising within the same culture.

The variation in OFX-R frequencies when selecting on the same concentration of OFX may be driven by several, non-exclusive biological factors. Firstly, the *Mtb* strains we tested may have different baseline DNA mutation rates. Secondly, the number and relative frequency of potential mutations that confer OFX-R may vary depending on the *Mtb* genetic background. Thirdly, the relative cost of OFX-R mutations may differ between *Mtb* genetic backgrounds. As the observed frequency of OFX-R in *Mtb* is likely the result from a combination of multiple factors, we took advantage of the fact that we had identified strains with a range of OFX-R frequencies. We selected three representative strains with significantly different OFX-R frequencies: N0157, N1283, and N0145. These strains had a high, mid-, and low frequency of OFX-R, respectively (Fig. 1A). We then explored the relative contributions of each biological factor listed above in driving the variation in OFX-R across genetically distinct *Mtb* strains.

### Mutation rate differences do not drive the *in vitro* variation in ofloxacin-resistance frequency in *M. tuberculosis*

We first tested for the presence of differential mutation rates between our panel of *Mtb* strains in Fig. 1A. Mutations in *dnaE*, which encodes the replicative DNA polymerase and serves as the major replicative exonuclease in *Mtb*, have been shown to confer a hypermutator phenotype in *Mtb* in the absence of environmental stress (50, 51). While *dnaE* mutations were present in the genomic data of our panel of drug-susceptible *Mtb* strains (See SI Appendix, Table S2) (49), none were in the polymerase and histidinol phosphatase domain of DnaE, the region where mutations would impart a hypermutator phenotype (50, 51). We did not test for the presence of *dnaE* mutations in the resistant colonies following the fluctuation analysis, as we reasoned that the likelihood of gaining both a *dnaE* and a *gyrA* double mutation within this relatively short period is extremely low as to be considered negligible. To test for mutation rate variation *in vitro*, we again conducted a fluctuation analysis on N0157, N1283, and N0145 (the high-, mid-, and low-frequency OFX-R strains, respectively), but used streptomycin (STR; 100 µg/mL) instead of OFX. We hypothesized that if the frequency of OFX-R is driven by differential mutation rates, then we should expect similar differences in the frequency of STR-resistance (STR-R). However, we observed no evidence for differences in the frequency of STR-R between the strains tested (Fig. 2, *P* = 0.135, Kruskal-Wallis; See SI Appendix, Table S3). This suggested that the observed differences in frequency of resistance are specific to OFX, and that there are limited, if any, inherent differences in mutation rate between the *Mtb* strains tested.

**Fig. 2.**
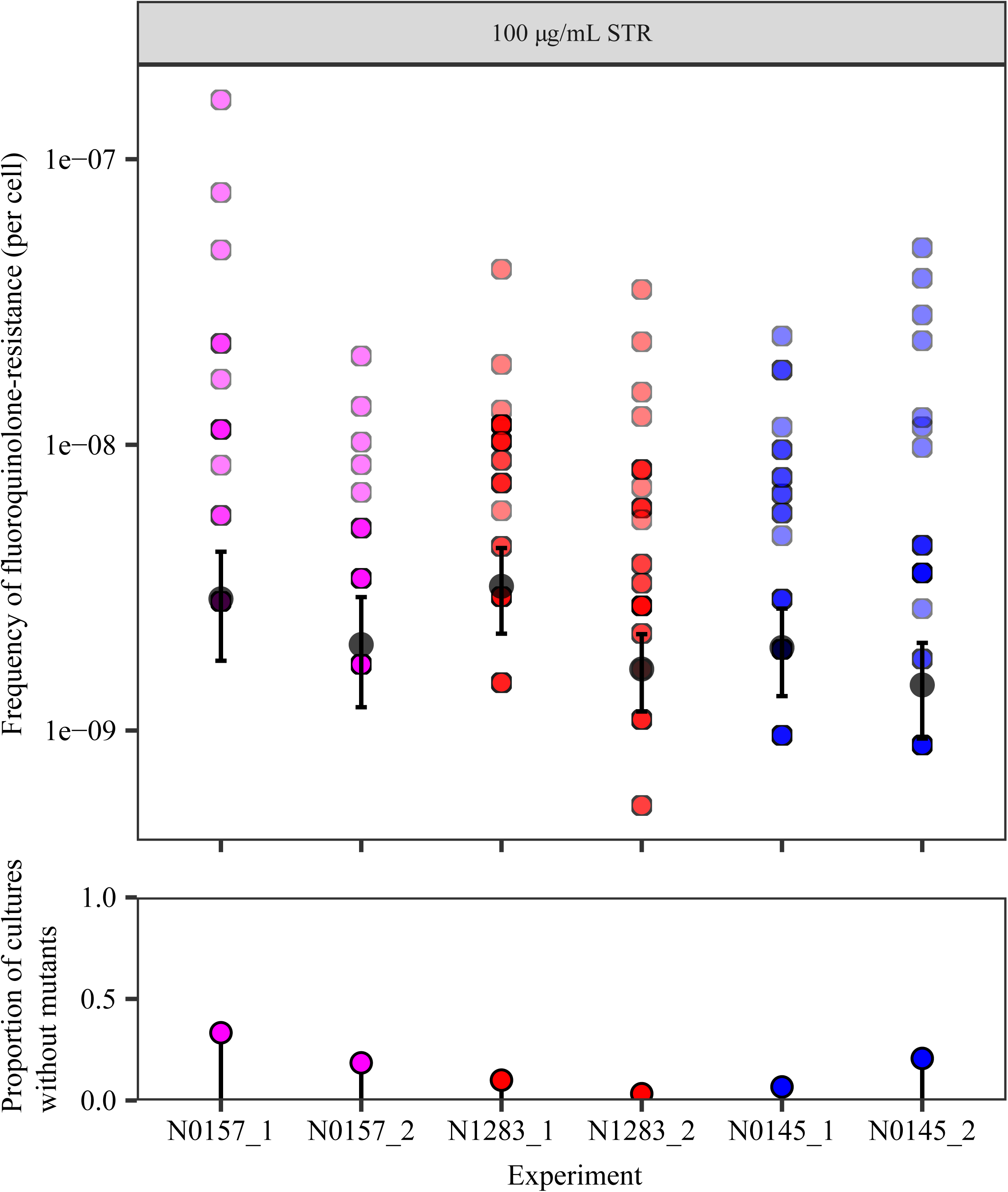
Frequency of streptomycin-resistance at 100 µg/mL streptomycin (STR) for wild-type N0157, N1283, and N0145 *M. tuberculosis* strains, as measured by fluctuation analysis assay. Top panel: Coloured points represent the frequency of resistant mutants per cell per parallel culture, with darker points representing multiple cultures with the same frequency. Colours denote the lineage that the *M. tuberculosis* strain belongs to (L1 = pink; L2 = blue; L4 = red). Grey points represent the estimated number of mutations per cell per strain as calculated by MSS-MLE, while black bars denote the respective 95% confidence intervals. Bottom panel: the percentage of parallel cultures lacking STR-resistant mutants. Two biological replicates are presented for each *M. tuberculosis* strain, with each replicate identifier suffixed after the strain name.

### Mutational profile for ofloxacin-resistance is highly strain-dependent

We next determined the mutational profile for OFX-R for each strain used in the fluctuation analysis at 4 µg/mL OFX (Fig. 1A). The QRDR mutations in 680 *gyrA* and 590 *gyrB* sequences were identified in the resistant colonies. We observed that *gyrA* mutations made up 99.7% of the QRDR mutations observed (645 *gyrA* mutations, 2 *gyrB* mutations), and no QRDR double-mutants were present (See SI Appendix, Tables S4-S5). The mutational profiles for OFX-R were also highly strain-specific (Fig. 3A, *P* = 5.00 × 10^-4^, Fisher’s exact test). Specifically, the GyrA A90V mutation was most prevalent in the high-frequency OFX-R strains, while GyrA D94G was most prevalent in all other strains. There was also a slight trend showing that strains with a greater number of unique *gyrA* mutations present also had higher rates of OFX-R (Fig. 1A; Fig. 3B).

**Fig. 3.**
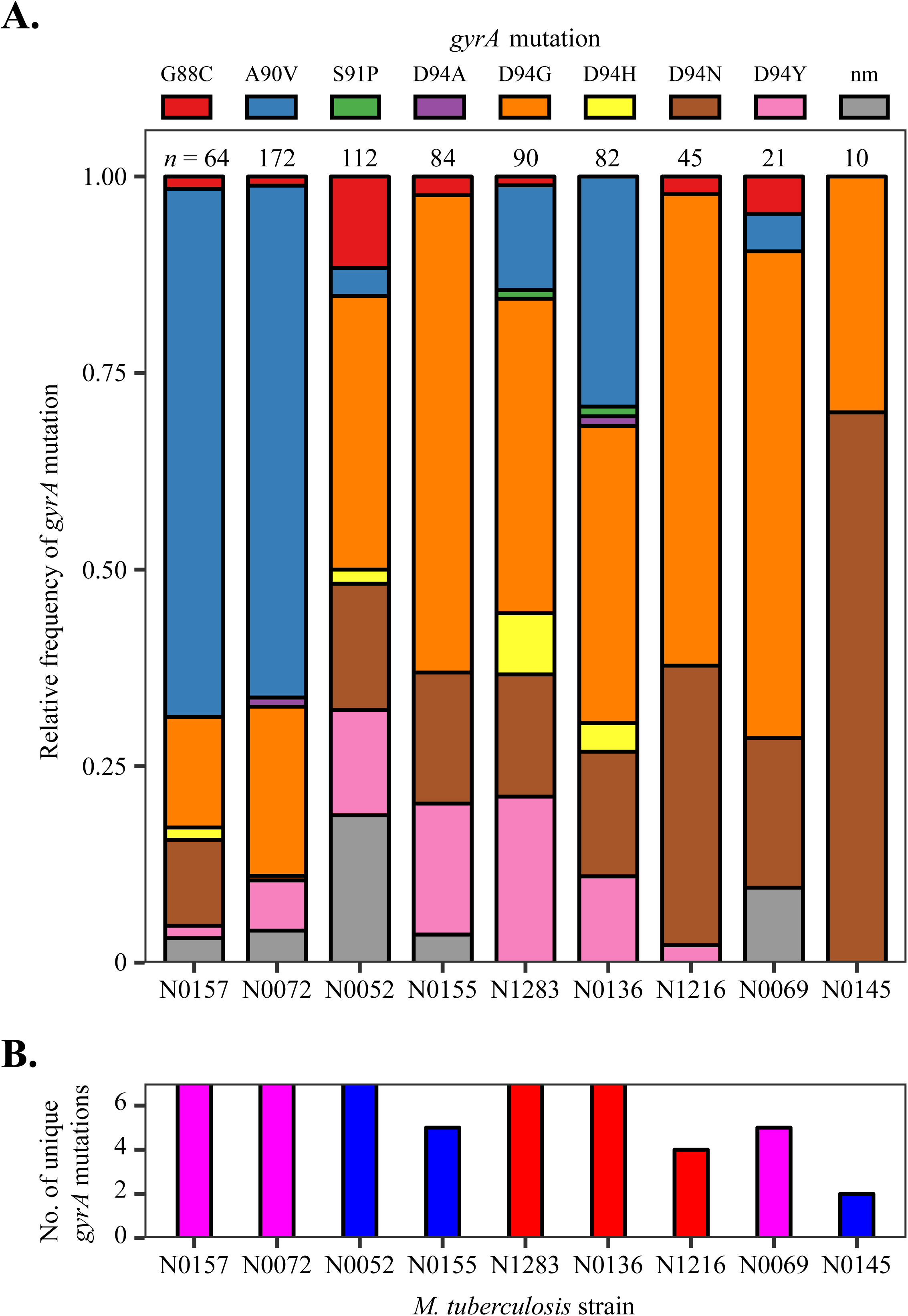
Variation in the mutational profile for ofloxacin-resistance after fluctuation analyses using nine genetically-distinct *M. tuberculosis* strains. **A.** Mutations in the quinolone-resistance-determining region (QRDR) of *gyrA* was analyzed in 680 ofloxacin (OFX)-resistant colonies from the fluctuation analysis performed in Fig. 2A (nm = no identified QRDR *gyrA* mutations). Strains are ordered left to right based on their frequency of OFX-resistance at 4 µg/mL OFX. Numbers of colonies analyzed per strain are reported directly above each column. **B**. The number of unique QRDR *gyrA* mutations per *M. tuberculosis* strain for OFX-resistance. Bar colours denote the *M. tuberculosis* lineage the strain belongs to (L1 = pink; L2 = blue; L4 = red).

The strain-dependent variation the mutational profile for OFX-R may be due to *gyrA* mutations conferring different resistance levels depending on the *Mtb* strain they are present in. To test this hypothesis, we first isolated OFX-R mutants carrying one of four possible GyrA mutations (G88C, A90V, D94G, or D94N) in the three strains used in Fig. 2: N0157, N1283, and N0145. The OFX MIC was determined for each of the twelve OFX-R mutant strains, along with their respective wild-type ancestors. We found that each parental wild-type strain had different susceptibilities to OFX, with N0157, N1283, and N0145 having OFX MICs of 2 µg/mL, 0.6 µg/mL, and 0.5 µg/mL, respectively (Fig. 4A; See SI Appendix, Table S6). This was consistent with the fluctuation analysis results shown in Fig. 1B. Furthermore, we observed that the OFX MIC conferred by a given *gyrA* mutation varied depending on the strain it was present in (Fig. 4B; See SI Appendix, Table S6). For example, mutants in the N0157 strain generally had higher OFX MICs than mutants in either the N0145 or N1283 strains. The only mutation that deviated from this trend was GyrA G88C, which conferred a higher OFX MIC when in the N0145 strain. Notably, the GyrA A90V mutation conferred a resistance level equal to or greater than 4 µg/mL OFX in the N0157 and N1283 strains, but not in N0145. This was consistent with the presence of GyrA A90V in the OFX-R mutational profile for N0157 and N1283, but not in N0145, in the fluctuation analysis using 4 µg/mL OFX (Fig. 1A; Fig. 3). In summary, the differences in OFX MIC reflected the strain-dependent mutational profiles for OFX-R in *Mtb*, as expected.

**Fig. 4.**
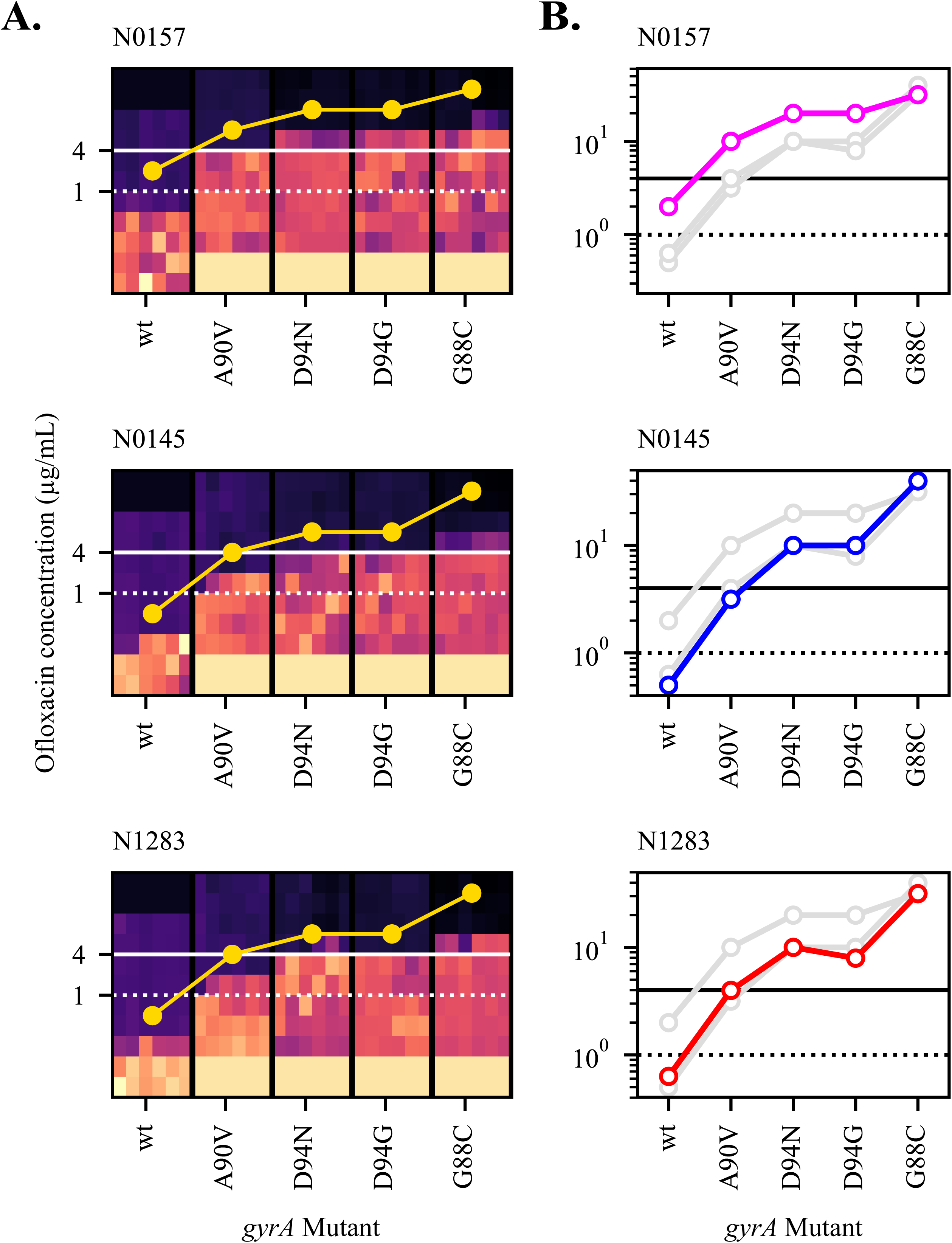
Ofloxacin (OFX) MIC is modulated by the genetic background of *M. tuberculosis*. **A.** Heat-map of OFX-susceptibility via Alamar Blue assay for *gyrA* mutant strains of *M. tuberculosis*, as well as their wild-type ancestor, in three genetic backgrounds (N0157, N0145, or N1283). Light areas represent growing cultures, while dark areas represent non-growing cultures. Yellow points represent estimates for OFX MIC (≥95% reduction in fluorescence). Areas of solid black colours (at 16+ μg/ml OFX for wild-type) and solid yellow colours (at <0.125 μg/ml OFX for mutants) were not measured and coloured in for illustrative purposes. **B**. OFX MIC estimates for each strain per genetic background, superimposed. Coloured points and lines represent MIC measurements for highlighted genetic background, with the line colour denoting the lineage that the strain belongs to (L1 = pink, L2 = blue, L4 = red). Grey points and lines represent the other two genetic backgrounds.

### Fitness of ofloxacin-resistance mutations are associated with their relative frequency *in vitro*

While the OFX MICs may determine which mutations may be observed in a fluctuation analysis, it is not the sole parameter to influence the OFX-R mutational profile for a given strain. We found that while the same *gyrA* mutation can be observed in two different *Mtb* strains, their relative frequencies may vary (Fig. 3). This variation may be due to the fitness of a given *gyrA* mutant being different across genetic backgrounds. To test this hypothesis, we used cell growth assays in antibiotic-free conditions to measure the *in vitro* fitness of our panel of OFX-R mutants relative to their respective parental wild-type ancestors. We observed that the relative fitness of the OFX-R mutants was modulated by both the *gyrA* mutation and the *Mtb* strain they were present in (Fig. 5A; See SI Appendix, Fig. S2-S3, Table S7). Furthermore, there was a positive association between the fitness of a given *gyrA* mutation with its relative frequency in the fluctuation analysis for the N0157 and N1283 strains (Fig. 5B, *P* = 0.03 for N0157, *P* = 0.05 for N1283). There was no evidence of an association in the N0145 background due to the lack of GyrA G88C and A90V mutants in its fluctuation analysis.

**Fig. 5.**
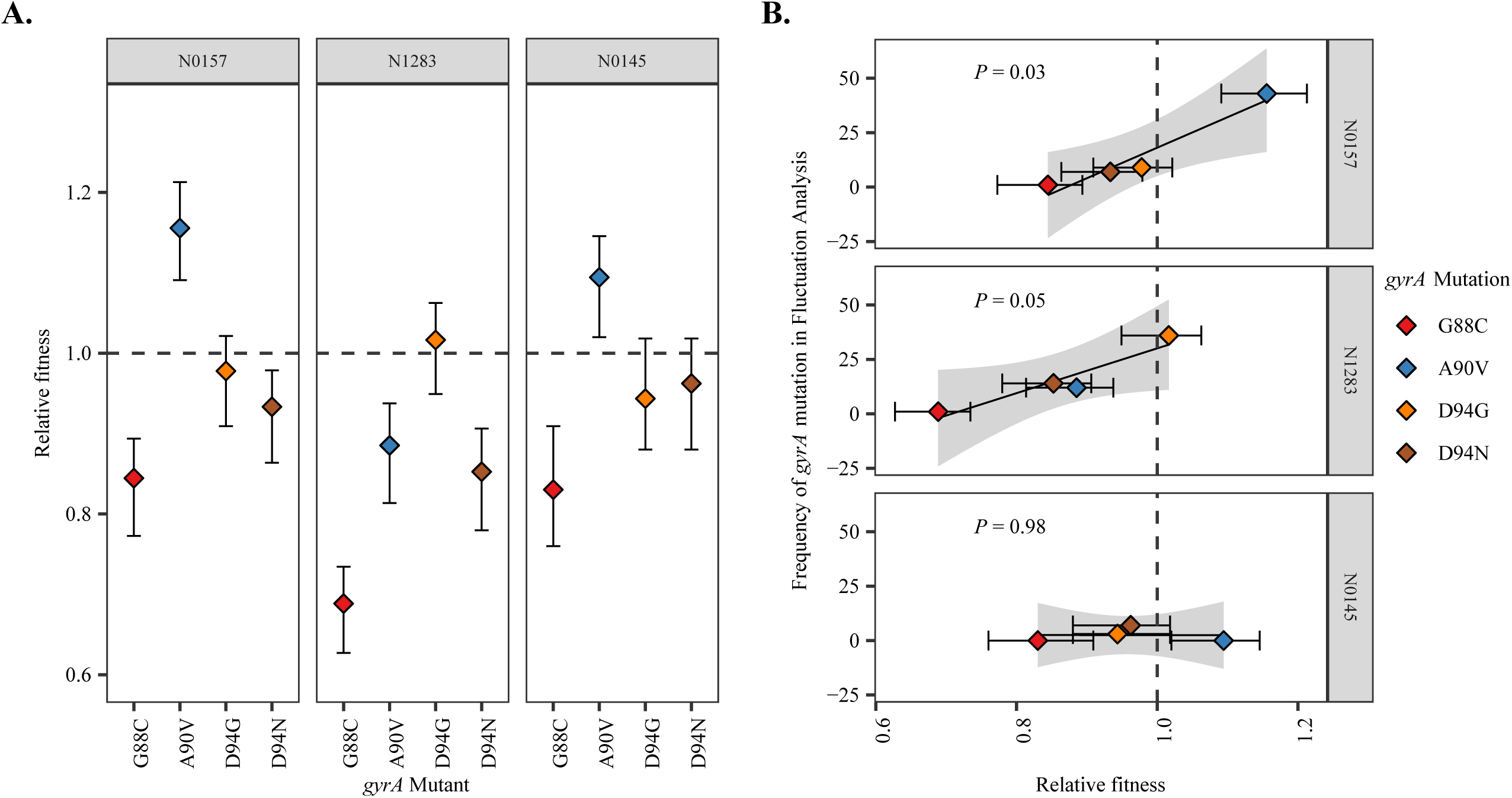
The *M. tuberculosis* genetic background modulates the fitness effect of fluoroquinolone-resistance mutations. **A.** Fitness of ofloxacin-resistant *M. tuberculosis* strain with specified *gyrA* mutation relative to the fitness of their respective wild-type ancestral strain. Ancestral strain per *gyrA* mutant is indicated in the grey bar above each panel. Fitness was measured by cell growth assay in antibiotic-free conditions. **B**. Association between the relative fitness of specified *gyrA* mutant and their absolute frequency during the fluctuation analysis performed in Fig. 1A, in three genetic backgrounds (N0157, N1283, and N0145).

The results from Fig. 4 and Fig. 5, as well as the apparent lack of mutation rate differences between our strains (Fig. 1C), suggested that differential mutational profiles was an important contributor in the variation in OFX-R frequency in *Mtb*. These mutational profile differences appear to be driven by the *Mtb* genetic background’s effect on both the MIC and the relative fitness cost of OFX-R mutations. We next explored whether these *in vitro* results would be relevant in clinical settings.

### Mutational profile for fluoroquinolone-resistance *in vitro* reflects clinical observations

To explore the clinical relevance of our *in vitro* work, we surveyed the FQ-R mutational profile from publicly available *Mtb* genomes obtained from clinical isolates. FQs are generally used for treatment against MDR-TB (29). While it is unclear whether resistance mutations for isoniazid (INH) and/or rifampicin (RIF) predispose a strain to become FQ-R, the prevalence of FQ-R is heavily biased towards MDR-TB strains due to treatment practices. We therefore based our analyses on a collated dataset of 3,452 publicly available MDR-TB genomes (See SI Appendix, Table S8), which we confirmed to be MDR-TB based on the presence of known INH- and RIF-resistance mutations. This dataset provided a reasonable sampling of the overall genetic diversity of *Mtb*, as six of the seven known phylogenetic *Mtb* lineages were represented (Lineages 1 – 6) (17, 30). We catalogued their FQ-R mutational profiles, and found 950 FQ-R mutations in 854 genomes (See SI Appendix, Tables S9-S10), showing that multiple FQ-R mutations may be present in the genome of a single *Mtb* clinical isolate. The frequency of FQ-R differed between lineages, with the highest frequencies present in L2 and L4 strains (*P* < 2.2 × 10^-16^, Chi-square Goodness of Fit Test). Moreover, we noticed a lineage-dependent mutational profile for FQ-R (Fig. 6, *P* = 3.00 × 10^-5^, Fisher’s exact test; See SI Appendix, Fig. S4, Tables S10-S11). For example, while the GyrA D94G mutation was most prevalent in strains belonging to L1, L2, and Lineage 3 (L3), the GyrA A90V mutation was most prevalent in L4 and Lineage 6 (L6).

**Fig. 6.**
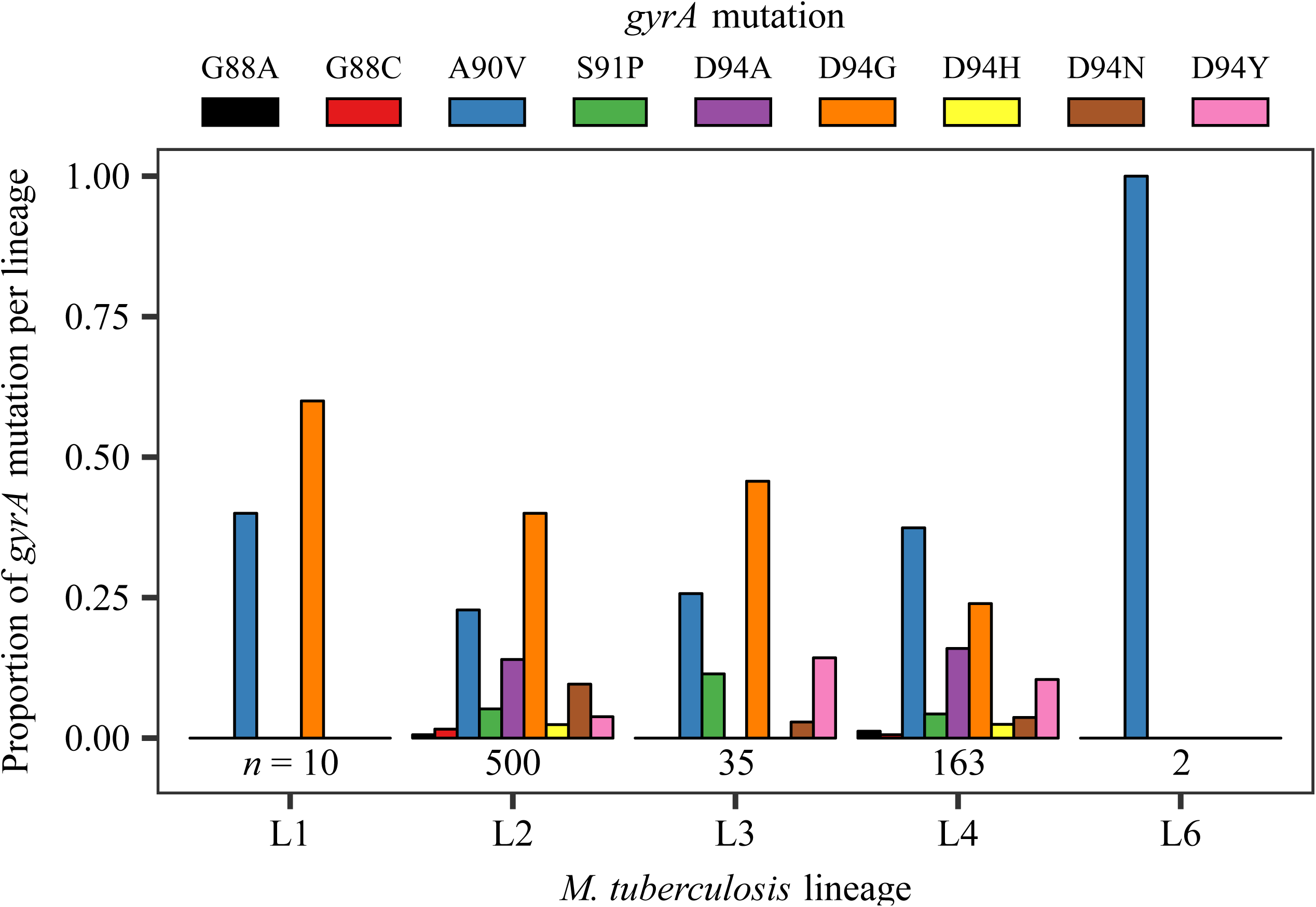
Mutational profile for fluoroquinolone-resistance *gyrA* mutations in clinical isolates of *M. tuberculosis*, per lineage. An initial dataset consisting of 3,452 genomes with confirmed MDR-TB mutations were surveyed. 854 genomes were identified as fluoroquinolone-resistant, with 848 of these genomes containing *gyrA* mutations. Only fixed fluoroquinolone-resistance mutations in the *gyrA* gene are enumerated here (*n* = 710). No fixed mutations were observed in L5 strains. Numbers of genomes analyzed per lineage are presented directly below their respective bar graph.

We observed that the mutational profile for FQ-R in the fluctuation analysis experiments mimicked published clinical data. Firstly, *gyrA* mutations made up the large majority of FQ-R mutations *in vitro* (Fig. 3; See SI Appendix, Tables S4-S5) and 944 out of the 950 QRDR mutations in the clinic (99.6%; Fig. 6; See SI Appendix, Table S10). The relative frequencies of *gyrA* mutations for each genetic background *in vitro* were also similar to their relative frequencies in the clinic. We compared the frequency of *gyrA* mutations from the OFX-R mutational profile assay in Fig. 3 to our genomic data survey in Fig. 6, but limited it to L2 and L4 strains (the two lineages with the highest clinical frequencies of FQ-R). We observed a positive association between the frequency of a given *gyrA* mutation in our fluctuation analysis compared to the frequency in the clinic, with the association being significant for L2 strains (Fig. 7, *P =* 0.027 for L2, *P* = 0.130 for L4, Fisher’s exact test). Based on the adjusted *R^2^* values, 45% of the variability in the clinical frequency of *gyrA* mutations in L2 strains and 19% of the variability in L4 strains can be attributed to how FQ-R evolves in *Mtb in vitro*. As the *in vitro* evolution of FQ-R is itself modulated by the *Mtb* genetic background, this provided evidence for the *Mtb* genetic background’s role in the evolution of FQ-R in the clinic.

**Fig. 7.**
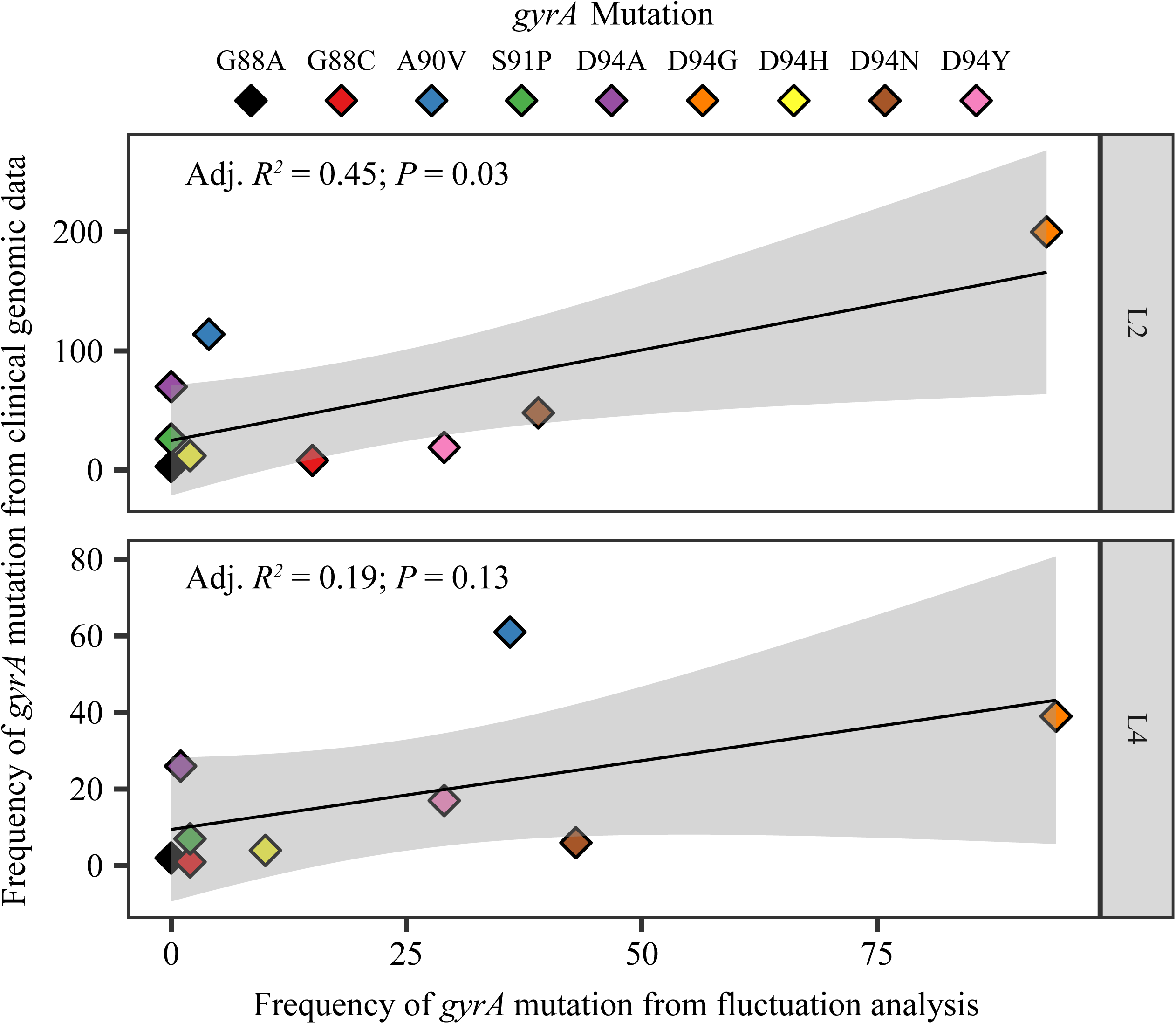
Association between the clinical frequencies of fluoroquinolone-resistance *gyrA* mutations with their respective *in vitro* frequencies amongst *M. tuberculosis* strains belonging to either the L2 or L4 lineages. Clinical frequencies are identical as reported in Fig. 6, while the *in vitro* frequencies are the same as in Fig. 3A, grouped by lineage.

## Discussion

Overall, we illustrate the *Mtb* genetic background’s considerable role in the evolution of resistance to FQs, a clinically important antimicrobial. We first explored whether the genetic variation among natural populations of *Mtb* can influence FQ-R evolution *in vitro*. Specifically, considering that *Mtb* treatment regimens are based on standardized antimicrobial concentrations (29), we tested whether different genetic variants of *Mtb* would acquire FQ-R at the same frequency when exposed to the same concentration of FQ. Fluctuation analysis on nine, genetically distinct, drug-susceptible *Mtb* strains showed that the genetic background can have a drastic effect on the rate of OFX-R acquisition when using the same concentration of OFX (Fig. 1). However, the effect of the genetic background on AMR frequencies observed here in the context of OFX-R differed from those reported in previous work focusing on other antibiotics. Specifically, experimental evidence from Ford *et al.* suggested that L2 Beijing strains have a higher basal DNA mutation rate compared to L4 (11), which consequently leads to a higher frequency of resistance against INH, RIF and ethambutol, even after correcting for differences in AMR mutational profiles. Based on these results, one would expect that L2 Beijing strains would also show higher frequencies of FQ-R. However, this was not the case in our fluctuation analysis for OFX-R, as one of our L2 Beijing strains (N0145) repeatedly acquired the lowest frequency of OFX-R (Fig. 1). Moreover, we saw minimal, if any, DNA base-pair mutation rate differences between three *Mtb* strains with different *in vitro* OFX-R frequencies (Fig. 2). Contradicting results on the *in vitro* frequency of AMR in *Mtb* have been reported before, with other fluctuation analyses showing no difference in the frequency of RIF-R emergence between L2 Beijing and non-L2 Beijing strains (52). Although diverging in their results, these previous studies, together with the study conducted here, highlight the importance of the genetic background when testing for the frequency of AMR in *Mtb*. Furthermore, these results show that differential DNA mutation rate is not the only parameter relevant in determining the frequency of FQ-R in *Mtb*.

If DNA mutation rates do not contribute to the variation in OFX-R frequency, we hypothesized that differences in the phenotypic effects of OFX-R mutations, and their consequent effect on the mutational profiles for OFX-R, may be important contributors. By sequencing the QRDR from resistant colonies in our OFX fluctuation analysis, we observed strain-specific patterns in the mutational profiles for OFX-R (Fig. 3). This suggested that the mutational profile for FQ-R is not only a function of the FQ type and concentration (10, 14, 47, 53), but that epistatic interactions between a given FQ-R mutation and the genetic background may also play a role. Similar epistatic interactions have been observed in *Escherichia coli* (26), *Pseudomonas* spp. (16, 27), *M. smegmatis* (54), and *Mtb* (24, 28, 31), where a given RIF-R *rpoB* mutation conferred differential MIC and fitness costs depending on the genetic background it occurred in, or on the presence of other AMR mutations. In line with these previous studies, we found that the OFX MIC and the fitness effect conferred by a given *gyrA* mutation varied significantly depending on the *Mtb* genetic background they occur in (Fig. 4; Fig. 5A; See SI Appendix, Table S6). These results support the hypothesis that epistasis plays a role in determining the strain-dependent OFX-R frequencies and mutational profiles observed during our fluctuation analyses (Fig. 3; Fig. 5B).

These epistatic interactions may have clinical consequences. A recent study has shown that drug-susceptible *Mtb* strains with higher MICs to INH and RIF were associated with increased risk of relapse following first-line treatment (20). Specific FQ-R *gyrA* mutations have also been associated with poorer treatment outcomes in MDR-TB patients (40, 55). Considering our observation that the *Mtb* genetic background affected both the OFX MICs and OFX-R mutational profiles (Fig. 3; Fig. 4; See SI Appendix, Tables S4-S6), the genetic background may therefore contribute to differences in patient treatment outcomes when using FQs as first-line drugs.

Using publicly available genomic data from *Mtb* clinical isolates, we observed significant lineage-dependent variation in the frequency of and mutational profiles for FQ-R (Fig. 6). As expected, the vast majority of FQ-R mutations were observed in *gyrA* (10, 22, 38, 39, 41–43). FQ-R was also most frequent in L2 and L4. This was also as expected, as strains from the L2 Beijing sublineage are known to associate with MDR-TB (4, 15, 21, 22), while L4 strains are the most prevalent globally, including in regions classified as high-burden for TB (17, 29, 56, 57). Consequently, strains from L2 and L4 would be more exposed to FQs, leading to the higher FQ-R frequencies observed in these two lineages. Furthermore, we observed that almost half of the variability in the clinical frequency of *gyrA* mutations of L2 strains can be explained by how *Mtb* evolves *in vitro* (Fig. 7). However, the *in vitro* FQ-R evolution could only account for 19% of the variability for *gyrA* mutation frequencies in clinical L4 strains. This suggested that while the *Mtb* genetic background can influence the evolution of FQ-R in the clinic, other factors (which may be independent of the *Mtb* genetic background) likely played strong roles as well. Epidemiological factors including socioeconomic disruptions, health system inefficiencies, and human behaviour are well known risk factors for the emergence and transmission of AMR in *Mtb* (3–7). Meanwhile, biological factors not explored in this study, such as antibiotic type and concentration (10–13, 46, 47), pharmacodynamic and pharmacokinetic features (58, 59), and the selective pressure of the host immune system (60), may also influence the evolution of FQ-R.

Our study is limited by the fact that our survey of clinical FQ-R frequencies involved a genomic dataset that was sampled by convenience. This dataset was used due to its public availability, and may not be fully representative of FQ-R frequencies in *Mtb* populations. We noted that lineage-specific frequencies of FQ-R were likely biased due to the overrepresentation L2 and L4 strains. Thus, to acquire a better understanding on which FQ-R mutations appeared and at what frequency they occurred at in different *Mtb* lineages, either more genomes from clinical isolates from other *Mtb* lineages must be made available, or a population-based study must be undertaken, preferably in a high burden MDR-TB region.

Exposure to quinolones have been shown to lead to SOS response-mediated mutagenesis, which can increase the rate of AMR acquisition, including resistance to quinolones themselves (53, 61, 62). Therefore, the strain-dependent OFX-R acquisition rates (Fig. 1) may be due to strain-dependent differences in the magnitude of quinolone-induced mutagenesis. We did not explicitly test for this possibility. However, phylogenetic SNPs present in SOS response-related genes may lead to strain-dependent differences in quinolone-induced mutagenesis, and we observed no such SNPs present across our panel of drug-susceptible *Mtb* strains (See SI Appendix, Table S2) (49). Thus, we observed no genetic evidence for strain-specific SOS response-mediated mutagenesis. Furthermore, in *E. coli*, quinolone-induced quinolone-resistant mutations may only be observed after 5 days of incubation with quinolones, which is equivalent to approximately 225 generations for wild-type *E. coli* (53, 61). Meanwhile, our wild-type *Mtb* strains were incubated for 40 generations at most in the presence of OFX (see Materials and Methods; See SI Appendix, Table S7), making the likelihood of observing OFX-induced OFX-R mutants in our *in vitro* system extremely low.

Another limitation of our study is that fluctuation analyses only model AMR emergence. Long-term population dynamics also play an important role in AMR evolution (8, 12, 14). For example, population bottleneck events modulate AMR evolution during serial transfer experiments (14, 27, 63, 64), and have also been hypothesized to strongly influence *Mtb* evolution in the clinic (65). Thus, modeling FQ-R evolution in *Mtb* in epidemiological settings would benefit from the use of some measure of long-term population dynamics and between-host transmission. Nevertheless, the fitness of AMR mutants is an important factor in determining its evolutionary fate (8, 9, 12, 14, 26, 54, 64) and its potential for between-host transmission (63, 66, 67). Considering that the *Mtb* genetic background modulated the fitness effect of FQ-R mutations (Fig. 5; See SI Appendix, Table S7), the genetic background may modulate how likely FQ-R mutants transmit between patients.

In conclusion, we illustrate how the genetic variation present in natural populations of *Mtb* modulates FQ-R evolution. Considering the non-random geographic distribution of different *Mtb* genetic variants (17, 30), our work suggests that there may be regional differences in the rate of FQ-R emergence and FQ-R prevalence when using FQs as a first-line drug. We therefore highlight the importance of bacterial genetics in determining how FQ-R evolves in *Mtb* and, in general, how AMR evolves in pathogens.

## Materials and Methods

### Collection of drug-susceptible clinical isolates of *M. tuberculosis* strains for *in vitro* studies

We used nine genetically-distinct *Mtb* strains, with three strains from each of the following *Mtb* lineages: Lineage 1 (L1; also known as the East-Africa and India Lineage), Lineage 2 (L2; the East Asian Lineage), and Lineage 4 (L4; the Euro-American Lineage) (17, 68). All strains were previously isolated from patients, fully drug-susceptible, and previously characterized by Borrell *et al*. (49) (See SI Appendix, Table S1).

Prior to all experimentation, starter cultures for each *Mtb* strain were prepared by recovering a 20 μL aliquot from frozen stocks into a 10 mL volume of Middlebrook 7H9 broth (BD), supplemented with an albumin (Fraction V, Roche), dextrose (Sigma-Aldrich), catalase (Sigma-Aldrich), and 0.05% Tween® 80 (AppliChem) (hereafter designated as 7H9 ADC). These starter cultures were incubated until their optical density at wavelength of 600 nm (OD_600_) was approximately 0.50, and were then used for *in vitro* assays.

### Fluctuation analyses

Fluctuation analyses were performed as described by Luria & Delbrück (48). Briefly, an aliquot from the starter cultures for each strain was used to inoculate 350 mL of 7H9 ADC to have an initial bacterial density of 5,000 colony forming units (CFU) per mL. This was immediately divided into 33 parallel cultures, each with 10 mL of culture volume aliquoted into individual 50 mL Falcon™ Conical Centrifuge Tubes (Corning Inc.). The parallel cultures were incubated at 37°C on standing racks, with re-suspension by vortexing (Bio Vortex V1, Biosan) every 24 hours. Cultures were grown until an OD_600_ of between 0.40 to 0.65. Once at this density, final cell counts (*N_t_*) from three randomly chosen parallel cultures were calculated by serial dilution and plating on Middlebrook 7H11 (BD), supplemented with oleic acid (AppliChem), albumin, and catalase (hereafter referred to as 7H11 OADC). To calculate the number of resistant colonies (*r*), the remaining 30 parallel cultures not used for *N_t_* determination were pelleted at 800 g for 10 min. at 4°C using the Allegra X-15R Benchtop Centrifuge (Beckmann Coulter). The supernatants were discarded, and the bacterial pellets re-suspended in 300 μL of 7H9 ADC. The re-suspensions were spread on 7H11 OADC plates supplemented with the relevant drug concentration (2, 4, or 8 µg/mL of ofloxacin, or 100 µg/mL streptomycin; Sigma). Resistant colonies were observed and enumerated after 21 to 35 days of incubation, depending on the *Mtb* strain. The estimated number of mutations per culture (*m*) was estimated from the distribution of frequency of drug-resistance per cell (*r_dist_*) using the Ma, Sarkar, Sandri-Maximum Likelihood Estimator method (MSS-MLE) (69). The frequency of drug-resistance acquired per cell (*F*) per strain was then calculated by dividing the calculated *m* values by their respective *N_t_* values. The 95% confidence intervals for each *F* were calculated as previously described by Rosche & Foster (69). Hypotheses testing for significant differences between the *r_dist_*between strains for the fluctuation analyses at 4 µg/mL of OFX (Fig. 1A) and at 100 µg/mL of STR (Fig. 1C) were performed using the Kruskal-Wallis test; significant differences in the *r_dist_* between strains in the fluctuation analyses at 2 and 8 µg/mL (Fig. 1B) were tested for using the Wilcoxon rank-sum test. Statistical analyses were performed using the R statistical software (70).

### Determining the mutational profile for ofloxacin-resistance *in vitro*

From the parallel cultures plated on 4 µg/mL of OFX (Fig. 1A), up to 120 resistant colonies per strain (at least 1 colony per plated parallel culture if colonies were present, to a maximum of 6) were transferred into 100 μL of sterile deionized H_2_O placed in Falcon® 96-well Clear Microplate (Corning Inc.). The bacterial suspensions were then heat-inactivated at 95°C for 1 h, and used as PCR templates to amplify the QRDR in *gyrA* and *gyrB* using primers designed by Feuerriegel *et al*. (71). PCR products were sent to Macrogen, Inc. or Microsynth AG for Sanger sequencing, and QRDR mutations were determined by aligning the PCR product sequences against the H37Rv reference sequence (72). Sequence alignments were performed using the Staden Package, while the amino acid substitutions identification were performed using the Molecular Evolutionary Genetics Analysis Version 6.0 package. Fisher’s exact test was used to test for significant differences between the strains’ mutational profiles for OFX-R. Data analyses were performed using the R statistical software (70).

### Isolation of spontaneous ofloxacin-resistant mutants

Spontaneous OFX-resistant mutants were isolated from strains belonging one of three genetic backgrounds: N0157 (L1, Manila sublineage; high frequency of OFX-R), N1283 (L4, Ural sublineage; mid-frequency of OFX-R), and N0145 (L2, Beijing sublineage; low frequency of OFX-R). To begin, we transferred 50 μL of starter cultures for each strain into separate culture tubes containing 10 mL of fresh 7H9 ADC. Cultures were incubated at 37°C until OD_600_ of approximately 0.80, and pelleted at 800 *g* for 5 min at 4°C. The supernatant was discarded, and the pellet re-suspended in 300 μL of 7H9 ADC. The re-suspension was plated on 7H11 OADC (BD) supplemented with 2 μg/mL of OFX, and incubated until resistant colonies appeared (approximately 14 to 21 days). Resistant colonies were picked and re-suspended in fresh 10 mL 7H9 ADC, and incubated at 37°C. Once the culture reached early stationary phase, two aliquots were prepared. The first aliquot was heat-inactivated at 95°C for 1 h, and the *gyrA* mutation identified by PCR and Sanger sequencing, as described in the mutational profile for OFX-R assay. If the first aliquot harboured one of four OFX-r *gyrA* mutations (GyrA^D94G^, GyrA^D94N^, GyrA^A90V^, or GyrA^G88C^), the second aliquot was stored in −80°C for future use.

Prior to further experimentation with the spontaneously OFX-R mutant strains, starter cultures were prepared in the same manner as for the drug-susceptible strains.

### Drug susceptibility assay

We determined the OFX-susceptibility levels of our spontaneous OFX-resistant mutants and their respective drug-susceptible ancestors by performing the colorimetric, microtiter plate-based Alamar Blue assay (73). Briefly, we used a Falcon® 96-well Clear Microplate, featuring a serial two-fold dilution of OFX. For drug-susceptible strains, a range of OFX concentration from 15 μg/mL to 0.058 μg/mL was used. Meanwhile, for OFX-resistant strains, a range of 60 μg/mL to 0.234 μg/mL was used. Each well was inoculated with a 10 μL volume of starter culture to have a final inoculum of approximately 5 × 10^6^ CFU/mL. The plates were incubated at 37°C for 10 days. Following incubation, 10 μL of Resazurin (Sigma) was added to each well, and the plates were incubated for another 24 h at 37°C. After this incubation period, plates were inactivated by adding 100 μL of 4% formaldehyde to every well. Measurement of fluorescence produced by viable cells was performed on SpectraMAX GeminiXPS Microplate Reader (Molecular Devices). The excitation wavelength was set at 536 nm, and the emission wavelength at 588 nm was measured. Minimum inhibitory concentration (MIC) for OFX was determined by first fitting a Hill curve to the distribution of fluorescence, and then defining the MIC as the lowest OFX concentration where the fitted Hill curve showed a ≥95% reduction in fluorescence. Two sets of experiments were performed for every strain, with three technical replicates per experiment. Analyses of MIC data were performed and figures created using the numpy, scipy, pandas and matplotlib modules for the Python programming language.

### Cell growth assay

We set up three or four 1,000 mL roller bottles with 90 mL of 7H9 ADC and 10 mL borosilicate beads. Each bottle was inoculated with a volume of starter cultures so that the initial bacterial density was at an OD_600_ of 5 × 10^-7^. The inoculated bottles were then placed in a roller incubator set to 37°C, and incubated for 12 to 18 days with continuous rolling. OD_600_ measurements were taken once or twice every 24 hours. Two independent experiments in either triplicates or quadruplicates were performed per strain.

We defined the exponential phase as the bacterial growth phase where we observed a log_2_-linear relationship between OD_600_ and time; specifically, we used a Pearson’s *R*^2^ value ≥ 0.98 as the threshold. The growth rate of a particular strain was then defined as the slope of the linear regression model. The relative fitness of a given spontaneous OFX-R mutant was defined by taking the growth rate of the OFX-resistant mutant strain and dividing it by the growth rate of its respective drug-susceptible ancestor. Linear regression models for the cell growth assays data were performed using the numpy, scipy, pandas and matplotlib modules for the Python programming language, as well as the R statistical software (70).

### Surveying the fluoroquinolone-resistance profile from publicly available *M. tuberculosis* genomes

We screened public databases to download global representatives of *Mtb* genomes, as described by Menardo *et al*. (74). We selected genomes that were classified as MDR-TB based on the presence of both isoniazid (INH)- and rifampicin (RIF)-resistance mutations. This provided a dataset of 3,452 genomes with confirmed MDR-TB; their accession numbers are reported in Table S8 (See SI Appendix). These MDR-TB genomes were then screened for the presence of FQ-resistance mutations, and we identified 854 genomes that were classified as FQ-R.

The INH-, RIF-, and FQ-resistance mutations used for screening are the same mutations used by Payne, Menardo *et al*. (75), and are listed in Table S12 (See SI Appendix). A drug-resistance mutation was defined as “fixed” in the population when it reached a frequency of ≥90%. Meanwhile, a drug-resistance mutation was considered “variable” in the population when its frequency was between 10% to 90%; thus, multiple drug-resistance mutations may be present in the genomic data from a single *Mtb* clinical isolate.

## Supporting information

Table S8 - Accession number of genomes from M. tuberculosis clinical isolates with confirmed MDR-TB mutations

Table S12 - List of high-confidence drug-resistance mutations used to determine drug-resistance mutational profiles of genomes from clinical isolates

## Acknowledgements

The authors would like to thank Sebastian Bonhoeffer, Daniel Angst, and Diarmaid Hughes for providing critical comments on the manuscript. Calculations were performed at sciCORE (http://scicore.unibas.ch/) scientific computing core facility at University of Basel. Library preparation and sequencing was carried out in the Genomics Facility Basel. This work was supported by the Swiss National Science Foundation (grants 310030_166687, IZRJZ3_164171, IZLSZ3_170834 and CRSII5_177163), the European Research Council (309540-EVODRTB) and SystemsX.ch.

