## Supplementary material for "The Evolution of Fluoroquinolone-Resistance in *Mycobacterium tuberculosis* is Modulated by the Genetic Background": SI Appendix

Socinstrasse 57, 4051 Basel, Switzerland

T: +41 61 284 6983

F: +41 61 284 8101

**This PDF file includes:**

Figs. S1 to S4

Tables S1 to S12

References for SI Appendix citations

### Supplementary Figures

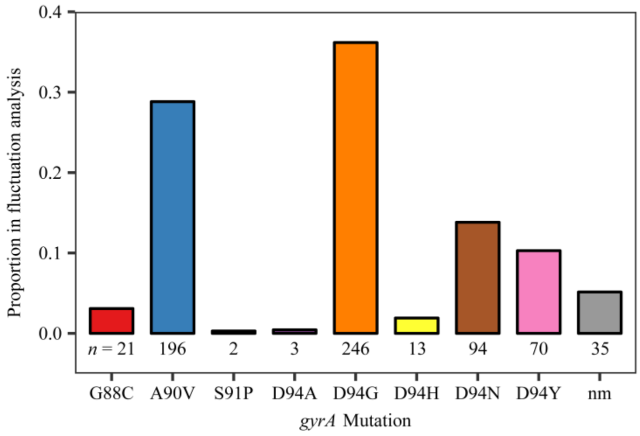


**Fig. S1**

Proportion of each *gyrA* mutation after sequencing of the QRDR of *gyrA* in 680 ofloxacin-resistant colonies from the fluctuation analysis performed in Fig. 2A (nm = no identified QRDR *gyrA* mutations). The numbers of colonies with the given *gyrA* mutation are reported directly below each respective column.

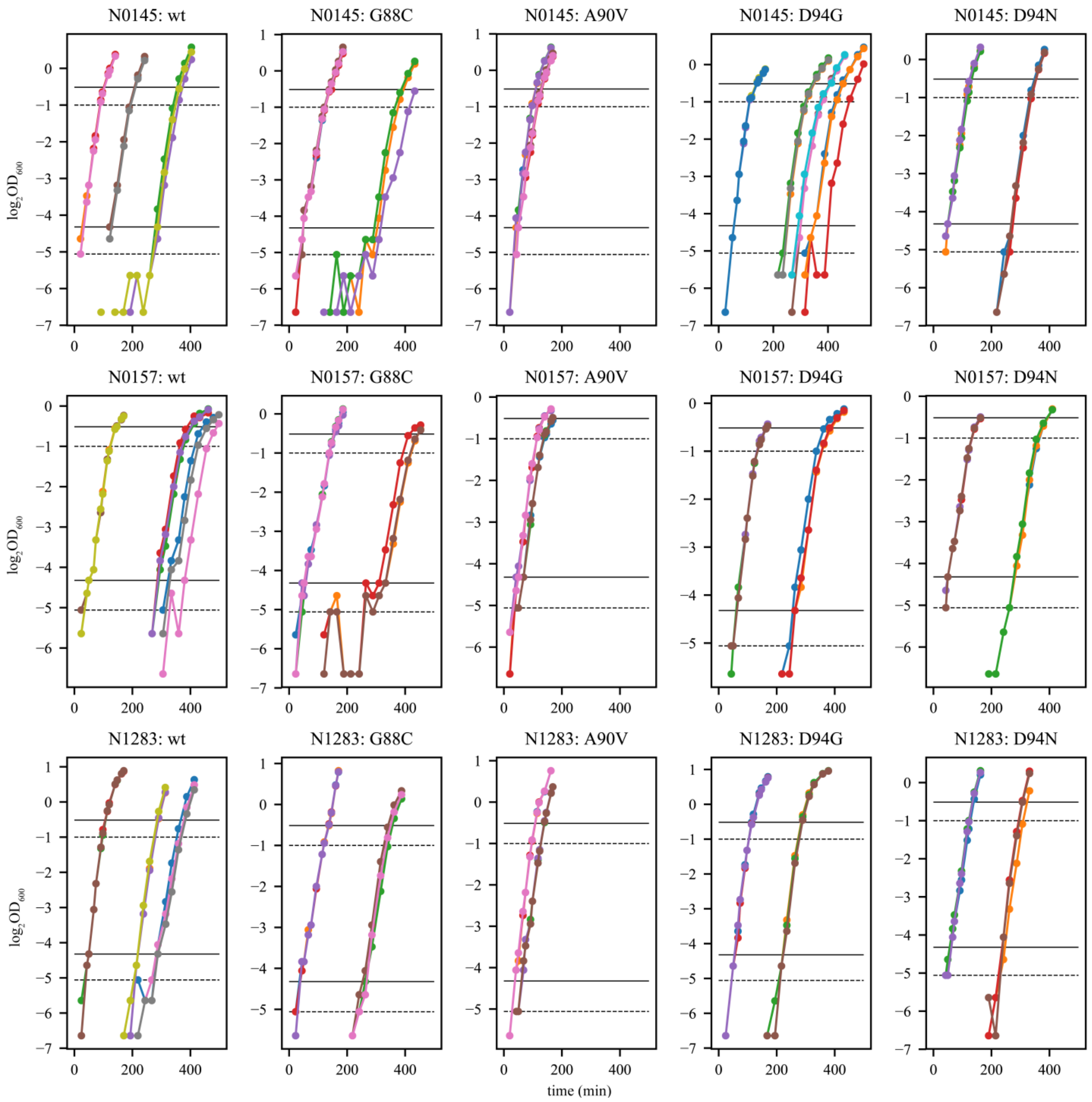


**Fig. S2**

Growth profiles of *M. tuberculosis* strains in cell growth assays under antibiotic free conditions, with all $OD_{600}$ values plotted ($log_2$-transformed). Genetic background of *M. tuberculosis* strain and its corresponding *gyrA* mutation were presented above each respective plot. Coloured dots represent the measured $log_2OD_{600}$ values at a given time (in minutes), with coloured lines connecting respective coloured dots. Different colours represent different replicates for each strain. For reference, black solid horizontal lines were plotted to denote non-transformed $OD_{600}$ values of either 0.05 (lower line) or 0.70 (upper line), while black dashed lines denote $OD_{600}$ values of either 0.03 (lower line) or 0.50 (upper line).

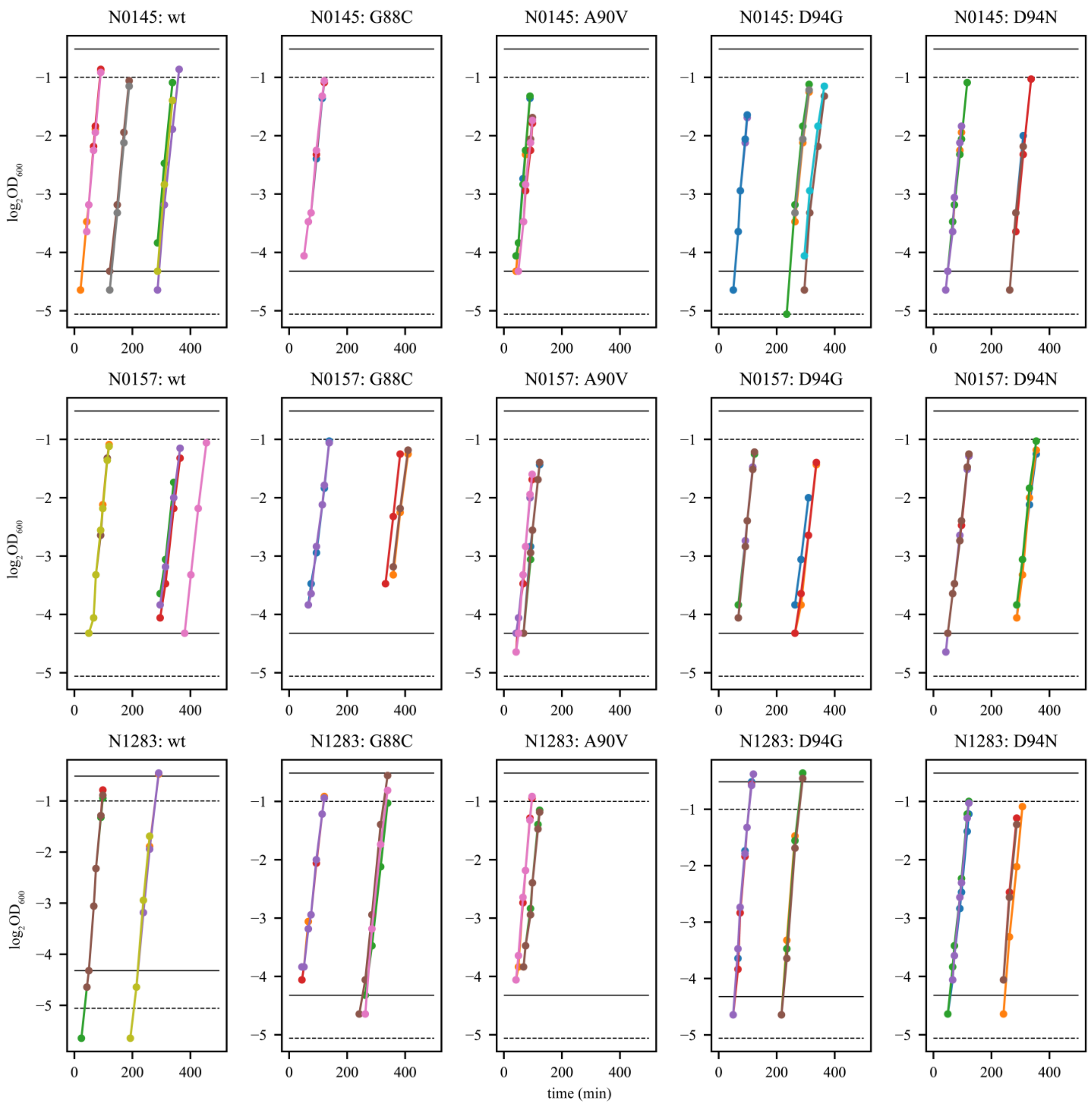


**Fig. S3**

Growth profiles of *M. tuberculosis* strains in cell growth assays under antibiotic free conditions, with only measured $OD_{600}$ values ($log_2$-transformed) present after filtering for exponential phase of growth. Genetic background of *M. tuberculosis* strain and its corresponding *gyrA* mutation were presented above each respective plot. Exponential growth phase was defined as a set consecutive time-points where a linear relationship between $\log_2 OD_{600}$ (defined by a Pearson's $R^2$ value ≥ 0.98) and time was present. Coloured dots represent the measured $\log_2 OD_{600}$ values at a given time (in minutes), with coloured lines connecting respective coloured dots. Different colours represent different replicates for each strain. For reference, black solid horizontal lines were plotted to denote non-transformed $OD_{600}$ values of either 0.05 (lower line) or 0.70 (upper line), while black dashed lines denote $OD_{600}$ values of either 0.03 (lower line) or 0.50 (upper line).

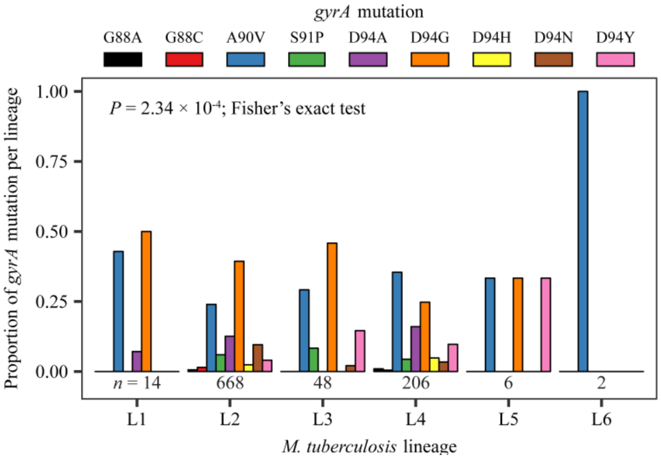


**Fig. S4**

Mutational profile for all (fixed and variable) fluoroquinolone-resistance *gyrA* mutations is lineage-specific in clinical isolates of *M. tuberculosis*. An initial dataset of 3,452 genomes with confirmed MDR-TB mutations were surveyed. 854 genomes were identified as fluoroquinolone-resistant, with 848 of these genomes containing *gyrA* mutations. If genomic data from a single *M. tuberculosis* clinical isolate contained multiple fluoroquinolone-resistance *gyrA* mutation, each mutation was counted once ($n$ = 944). Number of genomes analyzed per lineage is presented underneath each respective bar graph. *Mtb* lineage designations defined as in (1, 2).

### Supplementary Tables

**Table S1**

**Classification of *M. tuberculosis* strains used for *in vitro* assays**

| Strain | Lineage | Sub-lineage | Alternate Sub-lineage/Strain Nomenclature |
|---|---|---|---|
| N0069 | L1 | L1.1.1 | EAS042 (3) |
| N0072 | L1 | L1.1.2 | EAS053 (3) |
| N0157 | L1 | L1.2.1 | Manila; T92 (4) |

| | | | |
|---|---|---|---|
| N0052 | L2 | L2.2.2 | Beijing, Asia Central 1; 98_1833 (3, 5, 6) |
| N0145 | L2 | L2.2.1.1 | Beijing, Pacific RD150; T67 (5, 6) |
| N0155 | L2 | L2.2.1 | Beijing; T85 (4, 7) |
| N0136 | L4 | L4.3.3 | Latin America-Mediterranean; T4 (4, 5, 8) |
| N1216 | L4 | L4.6.2.2 | Cameroon (5, 8) |
| N1283 | L4 | L4.2.1 | Ural (5, 8) |

Lineage 1 = L1; Lineage 2 = L2; Lineage 4 = L4. *Mtb* lineage designations defined as in (1, 2).

**Table S2**

**Phylogenetic single nucleotide polymorphisms leading to missense mutations present in the genomic data of the nine dug-susceptible *M. tuberculosis* strains outlined in Supplementary Tables**

Table S1

| **Gene** | **Gene Name** | **Amino Acid Substitution** | **Strain** | **Lineage** |
|---|---|---|---|---|
| Rv0005 | *gyrB* | M291I | N0069 | L1 |
| Rv0005 | *gyrB* | M291I | N0072 | L1 |
| Rv0005 | *gyrB* | M291I | N0157 | L1 |
| Rv0006 | *gyrA* | A384V | N0069 | L1 |
| Rv0006 | *gyrA* | A384V | N0072 | L1 |
| Rv0006 | *gyrA* | A384V | N0157 | L1 |
| Rv0006 | *gyrA* | G247S | N0136 | L4 |
| Rv0006 | *gyrA* | K224E | N0072 | L1 |
| Rv0006 | *gyrA* | P154R | N1216 | L4 |
| Rv1547 | *dnaE1* | D316N | N0069 | L1 |
| Rv1547 | *dnaE1* | S898L | N0052 | L2 |
| Rv3370c | *dnaE2* | C313W | N0157 | L1 |
| Rv3370c | *dnaE2* | P814S | N0052 | L2 |

**Table S3**

**Mutations present in the *rpsL* gene for 194 streptomycin-resistant colonies following fluctuation analysis on 100 µg/mL of streptomycin**

| Strain | K43M | K43N | K43R | K43R, K10Q | K43T | K88E | K88R | nm |
|---|---|---|---|---|---|---|---|---|
| N0157 | 0 | 1 | 5 | 0 | 9 | 0 | 0 | 0 |
| N1283 | 4 | 4 | 72 | 1 | 13 | 3 | 5 | 9 |
| N0145 | 0 | 0 | 46 | 0 | 13 | 0 | 0 | 9 |

nm = streptomycin-resistant colonies with no mutations in *rpsL*

**Table S4**

**Mutations in the QRDR of *gyrA* for 680 ofloxacin-resistant colonies following fluctuation analysis on 4 µg/mL of ofloxacin**

| Strain | G88C | A90V | S91P | D94A | D94G | D94H | D94N | D94Y | nm | Strain Total |
|---|---|---|---|---|---|---|---|---|---|---|
| N0069 | 1 | 1 | 0 | 0 | 13 | 0 | 4 | 0 | 2 | 21 |
| N0072 | 2 | 112 | 0 | 2 | 37 | 0 | 1 | 11 | 7 | 172 |
| N0157 | 1 | 43 | 0 | 0 | 9 | 1 | 7 | 1 | 2 | 64 |
| N0052 | 13 | 4 | 0 | 0 | 39 | 2 | 18 | 15 | 21 | 112 |
| N0145 | 0 | 0 | 0 | 0 | 3 | 0 | 7 | 0 | 0 | 10 |
| N0155 | 2 | 0 | 0 | 0 | 51 | 0 | 14 | 14 | 3 | 84 |
| N0136 | 0 | 24 | 1 | 1 | 31 | 3 | 13 | 9 | 0 | 82 |
| N1216 | 1 | 0 | 0 | 0 | 27 | 0 | 16 | 1 | 0 | 45 |
| N1283 | 1 | 12 | 1 | 0 | 36 | 7 | 14 | 19 | 0 | 90 |
| **Mutation Total** | 21 | 196 | 2 | 3 | 246 | 13 | 94 | 70 | 35 | 680 |

nm = ofloxacin-resistant colonies with no mutations in the QRDR region of *gyrA*

**Table S5**

**Mutations in the QRDR of *gyrB* for 590 ofloxacin-resistant colonies following fluctuation analysis on 4 µg/mL of ofloxacin**

| Strain | E454K | D461H | nm | Strain Total |
|---|---:|---:|---:|---:|
| N0069 | 0 | 0 | 22 | 42 |
| N0072 | 0 | 0 | 155 | 155 |
| N0157 | 0 | 1 | 41 | 22 |
| N0052 | 1 | 0 | 101 | 102 |
| N0145 | 0 | 0 | 7 | 66 |
| N0155 | 0 | 0 | 66 | 7 |
| N0136 | 0 | 0 | 81 | 69 |
| N1216 | 0 | 0 | 46 | 81 |
| N1283 | 0 | 0 | 69 | 46 |
| **Mutation Total** | 1 | 1 | 588 | 590 |

nm = ofloxacin-resistant colonies with no mutations in the QRDR region of *gyrB*

**Table S6**

**Ofloxacin MIC estimates for *gyrA* mutant strains and their respective parental strain**

| **Strain** | ***gyrA* Mutation** | **Genetic Background (Parental Strain)** | **Ofloxacin MIC (µg/mL)** | **Normalized Ofloxacin MIC*** |
|---|---|---|---|---|
| N0157 | wt | --- | 2.00 | 1.00 |
| N3661 | G88C | *N0157* | 31.60 | 15.80 |
| N2034 | A90V | *N0157* | 10.00 | 5.00 |
| N2036 | D94G | *N0157* | 20.00 | 10.00 |
| N2035 | D94N | *N0157* | 20.00 | 10.00 |
| N1283 | wt | --- | 0.60 | 1.00 |
| N2508 | G88C | *N1283* | 12.60 | 21.00 |
| N2505 | A90V | *N1283* | 4.00 | 6.67 |
| N3915 | D94G | *N1283* | 12.60 | 21.00 |
| N2507 | D94N | *N1283* | 12.60 | 21.00 |
| N0145 | wt | --- | 0.50 | 1.00 |
| N3659 | G88C | *N0145* | 39.80 | 79.60 |
| N2847 | A90V | *N0145* | 3.20 | 6.40 |
| N1893 | D94G | *N0145* | 10.00 | 20.00 |
| N1895 | D94N | *N0145* | 10.00 | 20.00 |

MIC estimates for Ofloxacin based on fitting of a Hill curve to the distribution of fluorescence in an Alamar Blue assay (9). MIC is defined as the ofloxacin concentration where fitted Hill curve showed a ≥95% reduction in fluorescence. *Normalized Ofloxacin MIC is calculated by taking the ofloxacin MIC of a given *M. tuberculosis* strain and dividing it by the ofloxacin MIC of its respective wild-type parental strain; Normalized Ofloxacin MICs for each wild-type parental strain are therefore equal to 1.00.

**Table S7**

***In vitro* fitness of *M. tuberculosis* strains based on cell growth assays in antibiotic-free conditions**

| Strain | *gyrA* Mutation | Genetic Background (Parental Strain) | Growth Rate (GR) | GR: Lower 95% | GR: Upper 95% | Generation Time (in hours) | Relative Fitness (RF) | RF: Lower 95% | RF: Upper 95% | *P* |
|---|---|---|---|---|---|---|---|---|---|---|
| N0157 | wt | --- | 0.045 | 0.044 | 0.047 | 22.22 | | | | |
| N3661 | G88C | *N0157* | 0.038 | 0.034 | 0.042 | 26.36 | 0.844 | 0.773 | 0.894 | **<0.001*** |
| N2034 | A90V | *N0157* | 0.052 | 0.048 | 0.057 | 19.23 | 1.156 | 1.091 | 1.213 | **<0.001*** |
| N2036 | D94G | *N0157* | 0.044 | 0.04 | 0.048 | 22.73 | 0.978 | 0.909 | 1.021 | 0.354 |
| N2035 | D94N | *N0157* | 0.042 | 0.038 | 0.046 | 23.81 | 0.933 | 0.864 | 0.979 | **0.009*** |
| N1283 | wt | --- | 0.061 | 0.059 | 0.064 | 16.40 | | | | |
| N2508 | G88C | *N1283* | 0.042 | 0.037 | 0.047 | 23.81 | 0.689 | 0.627 | 0.734 | **<0.001*** |
| N2505 | A90V | *N1283* | 0.054 | 0.048 | 0.06 | 18.52 | 0.885 | 0.814 | 0.938 | **<0.001*** |
| N3915 | D94G | *N1283* | 0.062 | 0.056 | 0.068 | 16.13 | 1.016 | 0.949 | 1.062 | 0.638 |
| N2507 | D94N | *N1283* | 0.052 | 0.046 | 0.058 | 19.23 | 0.852 | 0.780 | 0.906 | **<0.001*** |
| N0145 | wt | --- | 0.053 | 0.05 | 0.055 | 18.87 | | | | |
| N3659 | G88C | *N0145* | 0.044 | 0.038 | 0.05 | 22.73 | 0.830 | 0.760 | 0.909 | **<0.001*** |
| N2847 | A90V | *N0145* | 0.058 | 0.051 | 0.063 | 17.24 | 1.094 | 1.020 | 1.145 | **0.016*** |
| N1893 | D94G | *N0145* | 0.05 | 0.044 | 0.056 | 20.00 | 0.943 | 0.880 | 1.018 | 0.107 |
| N1895 | D94N | *N0145* | 0.051 | 0.044 | 0.056 | 19.61 | 0.962 | 0.880 | 1.018 | 0.206 |

The growth rate of a particular *M. tuberculosis* strain was defined as the slope during exponential phase of bacterial growth, with the exponential phase of bacterial growth defined as where a $\log_2$-linear relationship existed between $OD_{600}$ and time using a Pearson's $R^2$ value $\geq 0.98$ as the threshold. Generation times were calculated by taking the inverse of the calculated growth rate. The relative fitness of a given *gyrA* mutant was defined by taking its growth rate and dividing it by the growth rate of its respective wild-type ancestor.

**Table S8**

**Accession number of genomes used from *M. tuberculosis* clinical isolates with confirmed MDR-TB mutations**

(attached as a separate CSV file due to excessive length; *n* = 3,452)

**Table S9**

**Number of publicly available genomes from *M. tuberculosis* clinical isolates used to survey the mutational profile for fluoroquinolone-resistance**

| ***Mtb* lineage** | **No. of MDR-TB Genomes** | **No. of MDR-TB + FQ-R Genomes** |
|---|---:|---:|
| Lineage 1 | 109 | 13 |
| Lineage 2 | 1,903 | 597 |
| Lineage 3 | 151 | 44 |
| Lineage 4 | 1,263 | 196 |
| Lineage 5 | 18 | 2 |
| Lineage 6 | 8 | 2 |
| **Total** | 3,452 | 854 |

*Mtb* = *M. tuberculosis*; MDR-TB = multidrug-resistant tuberculosis, defined as *Mtb* genomes that have both an isoniazid and a rifampicin-resistance mutation; FQ-R = fluoroquinolone-resistant, defined as *Mtb* genomes that have a fluoroquinolone-resistance mutation. Multiple drug-resistance mutations present in the genomic data from a single *Mtb* clinical isolate is possible (classified as "variable," and therefore not "fixed" for drug-resistance mutations); if a genome contained multiple drug-resistance mutations, then the genome is simply counted as MDR-TB or FQ-R once. *Mtb* lineage designations defined as in (1, 2)**.**

**Table S10**

**Frequency of all (fixed and variable) fluoroquinolone-resistance mutations from sample set of 3,452 MDR-TB genomes.**

| **Mutation** | **L1** | **L2** | **L3** | **L4** | **L5** | **L6** | **Mutation Total** |
|---|---|---|---|---|---|---|---|
| *gyrA* G88A | 0 | 4 | 0 | 2 | 0 | 0 | 6 |
| *gyrA* G88C | 0 | 10 | 0 | 1 | 0 | 0 | 11 |
| *gyrA* A90V | 6 | 160 | 14 | 73 | 2 | 2 | 257 |
| *gyrA* S91P | 0 | 40 | 4 | 9 | 0 | 0 | 53 |
| *gyrA* D94A | 1 | 84 | 0 | 33 | 0 | 0 | 118 |
| *gyrA* D94G | 7 | 263 | 22 | 51 | 2 | 0 | 345 |
| *gyrA* D94H | 0 | 16 | 0 | 10 | 0 | 0 | 26 |
| *gyrA* D94N | 0 | 64 | 1 | 7 | 0 | 0 | 72 |
| *gyrA* D94Y | 0 | 27 | 7 | 20 | 2 | 0 | 56 |
| *gyrB* D461N | 0 | 3 | 0 | 2 | 0 | 0 | 5 |
| *gyrB* N499D | 0 | 1 | 0 | 0 | 0 | 0 | 1 |
| **Lineage Total** | 14 | 672 | 48 | 208 | 6 | 2 | 950 |

If a genome was classified as "variable" for fluoroquinolone-resistance mutations, each *gyrA* or *gyrB* mutation present were counted once. Lineage 1 = L1, Lineage 2 = L2, Lineage 3 = L3, Lineage 4 = L4, Lineage 6 = L5, Lineage 6 = L6. *Mtb* lineage designations defined as in (1, 2).

**Table S11**

**Frequency of fixed fluoroquinolone-resistance mutations from sample set of 3,452 MDR-TB genomes.**

| **Mutation** | **L1** | **L2** | **L3** | **L4** | **L5** | **L6** | **Mutation Total** |
|---|---|---|---|---|---|---|---|
| *gyrA* G88A | 0 | 3 | 0 | 2 | 0 | 0 | 5 |
| *gyrA* G88C | 0 | 8 | 0 | 1 | 0 | 0 | 9 |
| *gyrA* A90V | 4 | 114 | 9 | 61 | 0 | 2 | 190 |
| *gyrA* S91P | 0 | 26 | 4 | 7 | 0 | 0 | 37 |
| *gyrA* D94A | 0 | 70 | 0 | 26 | 0 | 0 | 96 |
| *gyrA* D94G | 6 | 200 | 16 | 39 | 0 | 0 | 261 |
| *gyrA* D94H | 0 | 12 | 0 | 4 | 0 | 0 | 16 |
| *gyrA* D94N | 0 | 48 | 1 | 6 | 0 | 0 | 55 |
| *gyrA* D94Y | 0 | 19 | 5 | 17 | 0 | 0 | 41 |
| *gyrB* D461N | 0 | 3 | 0 | 2 | 0 | 0 | 5 |
| *gyrB* N499D | 0 | 1 | 0 | 0 | 0 | 0 | 1 |
| **Lineage Total** | 10 | 504 | 35 | 165 | 0 | 2 | 716 |

Only genomes classified as “fixed” for fluoroquinolone-resistance mutations were enumerated here. Notably, no “fixed” mutations were observed in Lineage 5 (L5) strains. Lineage 1 = L1, Lineage 2 = L2, Lineage 3 = L3, Lineage 4 = L4, Lineage 6 = L6. *Mtb* lineage designations defined as in (1, 2).

**Table S12**

**List of high-confidence drug-resistance mutations used to determine drug-resistance mutational profiles of genomes from clinical isolates of *M. tuberculosis***

(attached as a CSV file due to length; *n* = 67)
